# From cartoons to quantitative models in Golgi transport

**DOI:** 10.1101/2020.05.13.094383

**Authors:** D. Nicolas Quiros, Luis S. Mayorga

**Affiliations:** IHEM (Universidad Nacional de Cuyo, CONICET), Facultad de Ciencias Médicas, Facultad de Ciencias Exactas y Naturales, Mendoza, Argentina

## Abstract

Cell biology is evolving to become a more formal and quantitative science. In particular, several mathematical models have been proposed to address Golgi self-organization and protein and lipid transport. However, most scientific articles about the Golgi apparatus are still using static cartoons to represent their findings that miss the dynamism of this organelle. In this report, we show that schematic drawings of Golgi trafficking can be easily translated into an Agent-Based Model (ABM) using the Repast platform. The simulations generate an active interplay among cisternae and vesicles rendering quantitative predictions about Golgi stability and transport of soluble and membrane-associated cargoes. The models can incorporate complex networks of molecular interactions and chemical reactions by association with COPASI, a software that handles Ordinary Differential Equations. The strategy described provides a simple, flexible, and multiscale support to analyze Golgi transport. The simulations can be used to address issues directly linked to the mechanism of transport or as a way to incorporate the complexity of trafficking to other cellular processes that occur in dynamic organelles.

## INTRODUCTION

Intracellular trafficking is a fundamental process for eukaryotic cells. Macromolecules need to find their way along the endocytic and secretory pathways to their final destination in the interior of the cell or to be secreted to the extracellular medium. It is not easy to envision this active exchange of material between membrane-bound structures. As a rule, macromolecules do not leave the donor organelle to travel through the cytoplasm to be incorporated into the acceptor compartment. Hence, transport requires the interaction and exchange of soluble and membrane-associated components among closed compartments. Whether the exchange is direct between the compartments or mediated by tubulo-vesicular transport carriers, the process requires two opposite and complementary events. Membrane fusion that allows the mixing of two organelles, and membrane fission that mediates the segregation and sorting of molecules among the dividing structures.

The mechanism of membrane fusion has been studied in detail (Jahn and Fasshauer, 2012;Sudhof and Rothman, 2009;Wickner and Schekman, 2008). A central core of proteins is required for membrane apposition and bilayer destabilization to promote the opening and expansion of a fusion pore connecting the membrane-bound structures. Besides the protein complex required to overcome the energetic barrier involved in the formation of the pore, another set of proteins are needed to provide specificity to the process. Fusion must occur among compatible organelles to preserve the complex organization of the cell. As a rule, organelles surrounded by similar membrane domains have a higher probability of fusing.

Membrane fission is also a well-characterized process (Renard *et al*., 2018). Depending on their protein and lipid composition, different membrane domains bind membrane-deforming protein complexes, such as COPs, clathrin, sorting nexins, and others (Bonifacino and Glick, 2004) (McCullough *et al*., 2013). The deformations lead to the sorting of membrane domains by budding of tubules or vesicles that are now separated from the membrane of the original organelle.

The identity of membrane domains is not completely lost upon fusion. It is well recognized the presence of separate membrane domains within a single organelle (Miaczynska and Zerial, 2002). Membrane domains can then be considered as key building blocks of the cellular endomembrane system, and they have special characteristics for each subcellular compartment. Membrane domains are not static, and can dynamically change their composition. In this scenario, soluble and membrane-bound cargoes are directed to their final destination following the fusion/fission interplay among organelles. The final destination of a molecule depends on its behavior during these events. Soluble cargoes that do not interact with membranes are transported in the lumen of the organelles; hence, during fission, they are distributed according to the volume of the newly formed organelles. Fluid-phase cargoes are enriched in round, large-volume structures and excluded from small vesicles and tubules. Membrane cargoes with no particular affinity for a membrane domain, similar to soluble cargoes, distribute proportionally to the area of organelles. However, membrane cargoes frequently carry specific tags that interact with one or more adaptor proteins that strongly affect their destination during fission (Kim, 2016). In addition, lipids in membranes are organized in microdomains, and membrane-anchored factors are also recruited to specific lipid environments that are segregated during the formation of vesicles and tubules (Kumar *et al*., 2015).

Depict the detailed knowledge about the molecular mechanisms involved in transport and the large list of factors that have been identified, the underlying logic of the process is still not well understood. For example, a classical controversy between vesicular transport and maturation in Golgi transport is still present after several decades, and hundreds of experiments using very ingenious tools to discriminate between the two models. Interestingly, both are presented as possibilities in Cell Biology books (Alberts *et al*., 2015), reviews (Glick and Luini, 2011), and encyclopedias (Luini and Parashuraman, 2016). At present, the evidence points to maturation as the main transport mechanism (Glick and Luini, 2011). In yeast, where the Golgi is not organized in stacks, the maturation of a single cisterna containing a fluorescent cargo has been observed in real-time images (Kurokawa *et al*., 2019). Besides, mathematical modeling (Mani and Thattai, 2016) and experimental data about the mechanism of transport of Golgi resident proteins are consistent with maturation (Liu *et al*., 2018).

Part of the problem is that hypotheses in intracellular transport are in general qualitative, like most ideas in Cell Biology. They are presented as schematic representations of compartments connected by arrows and seldom translated to formal models with quantitative predictions. The dynamic nature of organelles that change position, shape, and composition makes difficult to develop simple formal models for intracellular transport. Our group has shown that modeling of a simplified endocytic route composed by early, sorting, recycling, and late endosomes is possible using two complementary techniques: i) Agent-Based Modeling (ABM) to handle movement, fusion, and fission of organelles, and ii) Ordinary Differential Equations (ODE), to deal with molecule interactions and chemical reactions. Simulations generated with this model accurately reproduce several experimental results and could be used as a platform to represent complex molecular events such as Rab conversion, endosomal acidification, transport of lysosomal enzymes, and hydrolysis of glycolipids (Mayorga *et al*., 2018).

The flexibility of ABM allows building simulations with quantitative outputs from schematic representations of biological processes. An agent can be anything from a single molecule to a complete organelle. The behavior of the agents can be specified with simple rules that parallel the way of thinking in informal models. For example, the fusion among two organelles can be specified as “If there are two structures close enough, and if their membrane domains are compatible, fuse them to form a single structure having the volume, area and membrane-associated and soluble components of the original organelles".

The goal of the present report is to show that the cartoons used to represent Golgi trafficking can be translated into ABM models that produce quantitative predictions about Golgi stability and the transport of soluble and membrane-associated cargoes.

## RESULTS

### Brief description of an ABM model for intracellular transport

In the ABM model for intracellular transport developed previously, organelles are membrane-bound structures, characterized by volume, area, and movement. The area of the organelles corresponds to the surrounding surface and it is covered by one or more membrane domains. The movement is determined by position, speed, and direction in a 2D space. Other agents included are microtubules. Organelles can move randomly or following the plus (to the plasma membrane) or minus (to the nucleus) direction when near microtubules. In ABM, each agent is interrogated about performing or not “actions", according to the specific situation of the agent. After all agents have performed or not the actions, the process is iterated with the new situation of the agents. In ABM, each iteration is called a “tick” and it is a variable that can be calibrated to represent physical time. For Golgi transport, the actions implemented where “movement", “fusion", “fission", “maturation", “influx", and “outflux". The frequency of each action can be specified by “ticks” (for example, do “maturation” every 3000 ticks) or assigning a probability. Actions with higher probability occur faster than action with low probability (Table I).

**Table I.** Actions frequency for the different models. Frequency are expressed as a fix number of ticks or as the probability of occurring in one tick

| actions\models | Maturation | Vesicular | Distillation |
| --- | --- | --- | --- |
| move | 1 | 1 | 1 |
| fusion | 0.001 | 0.006 | 0.006 |
| fission | 0.002, 0.01, or 0.1 | 0.12 | 0.5 |
| influx | fixed (every 3000 ticks) | 0.02 | 0.001 |
| outflux | fixed (every 3000 ticks) | 0.12 | 0.006 |
| maturation | fixed (every 3000 ticks) | 0 | 0 |

In the present model, the relatively static position of the Golgi apparatus was mimicked by restricting the movement of cisternae to a perinuclear position. In contrast, vesicles moved randomly unless they were near a microtubule; in this case, they moved toward the nucleus.

To fuse, two organelles should be in close proximity. The fusion probability depended on the compatibility of the membrane domains of the two interacting organelles. During fission, a vesicle or cisterna was formed carrying a single membrane domain. Fusion and fission preserved the area, volume and membrane domains of the organelles. In contrast, during maturation, all the membrane domains of an organelle were switched to (mature to) a single domain.

During influx, a new membrane domain was incorporated into the system by adding a new vesicle or cisterna, or by incorporating the domain to an existing cisterna. During outflux, a vesicle or cisterna was deleted from the model. The contents of the deleted organelle were summed to account for the transfer to a post-Golgi compartment.

Within these organelles, which dynamically change with time, soluble and membrane-associated cargo were included. The final destination of a cargo depended on its behavior during fission. Soluble cargoes distributed according to the volume, and membrane-bound cargoes according to the area of the dividing organelles. Some cargoes had affinity for a membrane domain, and during fission, they followed the distribution of this domain. Large cargoes could not be included in newly formed vesicles and were retained in large cisternae.

### Maturation Model

In its simplest formulation, the Maturation Model proposes that new cisternae are assembled in the cis side of the Golgi and that they mature to medial and trans cisternae with time. Finally, the cisternae leave the Golgi to become TGN structures. To retain Golgi-resident factors (such as glycosyltransferases), these proteins are recruited in vesicles that fuse with the upcoming cisterna and they became engaged in a cycle of maturation and retrograde transport. These features of the model are represented in the schematic drawing shown in Fig. 1A (modified from (Alberts *et al*., 2015)). According to this representation of Golgi transport, the process can be described by the following rules:

**Figure 1.**
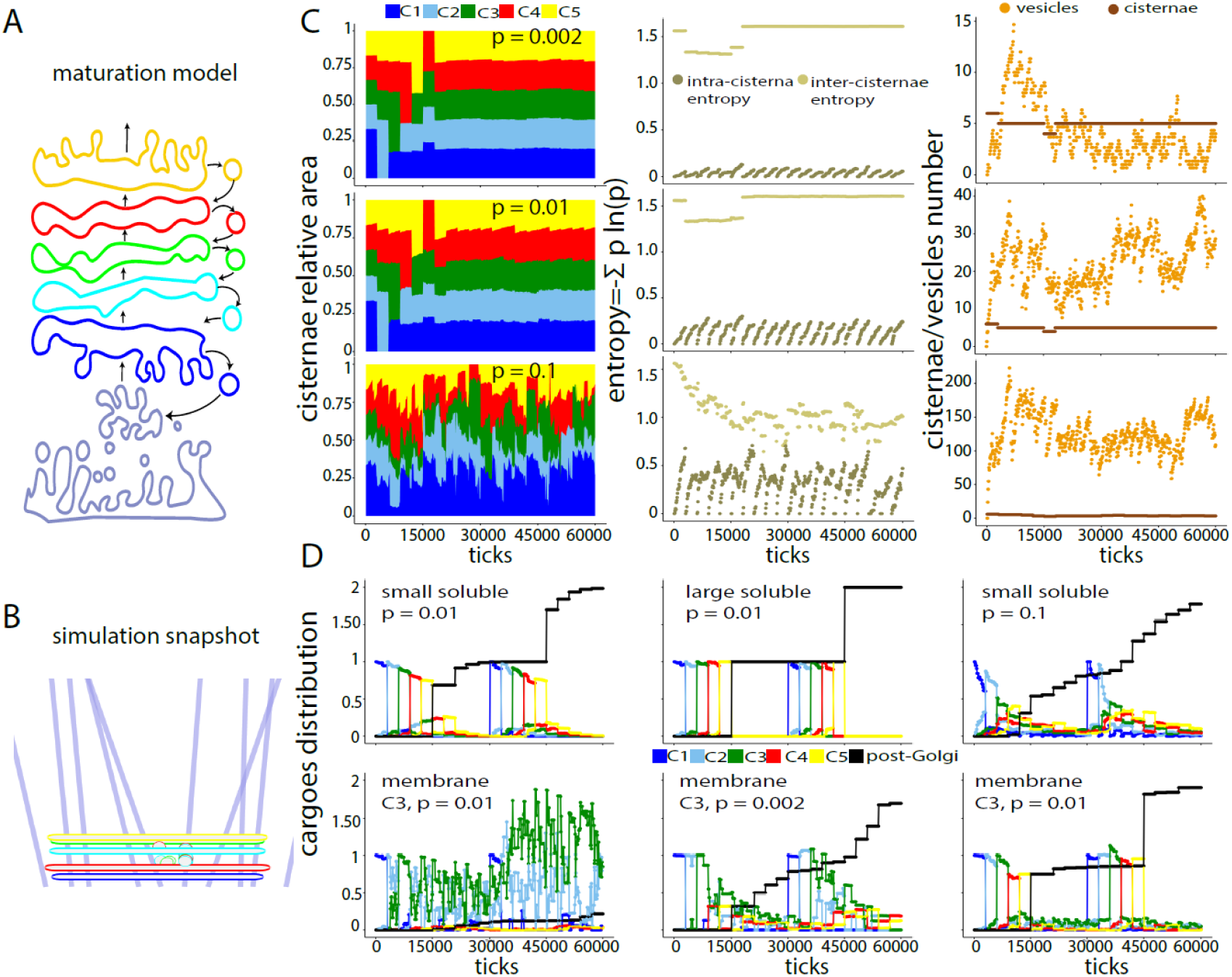
Maturation Model. **A**. Cartoon representing the maturation model. Notice the forward membrane flux/maturation process and the vesicle-mediated backward transport. **B**. Snapshot of the simulation built from the cartoon in A. **C**. Golgi stability when the probability of budding vesicles was increased from 0.002 to 0.01, or to 0.1. The parameters along the simulation shown are: the proportional cisterna area for each Golgi domain (left panels), the inter-cisternae and intra-cisterna entropies (central panels), and the number of vesicles and cisternae in the simulation (right panels). **D**. Transport of several cargoes under different vesicle budding probability (p). The cargoes (0.004 mM) were loaded at tick 1 and 30.000. The association of a small and a large soluble cargo with the different Golgi domains or a post-Golgi compartment was followed throughout the simulation (top panels). The large cargo could not fit into vesicles and was retained in cisternae. A fraction of the small cargo was transported backward in vesicles delaying its exit from the Golgi. In the bottom panels, the same association was plotted for a membrane-bound cargo that was allowed to enter into vesicles only in the C3 compartment. Notice that the cargo was efficiently retained in the C3 compartment (left panel). In the middle panel, very few vesicles were formed and the cargo left the Golgi. In the right panel, the amount loaded was increased to 0.4 mM and the vesicles formed were not sufficient to retain the cargo in the Golgi. In D, the results are normalized considering the maximal amount of cargo present in the simulation.

- New cisternae are formed in the cis side of the Golgi and are converted into TGN structures in the trans side of the organelle.
- The cisternae mature from cis to trans.
- Vesicles are formed in the cisternae carrying Golgi resident molecules and fuse to the previous cisterna to prevent the transport to the TGN.
- Vesicles are not allowed to fuse among them.
- Cisternae are not allowed to fuse among them.

To implement these rules in the ABM model, five “agents” with the characteristics of cisternae (500 nm in radius, 20 nm thick cylinders) were generated, each one carrying a different membrane domain (C1 to C5). The cisternae could form vesicles (fission) with the area and volume of a Cop I-type of structure (35 nm radius spheres). The vesicles could fuse with cisternae carrying a membrane domain corresponding to the previous cisterna in the cis-to-trans Golgi organization. For example, vesicles forming from C4 fused with the C3 cisterna (see fusion probability, Table II). Vesicles forming from the C1 cisterna were deleted (they were supposed to leave the Golgi to fuse with ER/ERGIC structures). Every 3000 ticks, a new C1 cisterna was added. Simultaneously, the old cisternae matured. This means that the old C1 cisterna switched to C2, C2 to C3, and so on. The C5 cisterna disappeared and the content was delivered to a post-Golgi compartment. A snapshot of the simulation is shown in Fig. 1B.

**Table II.** Fusion probabilities between Golgi domains for the three transport models

| maturation transport |  |  |  |  |  |
| --- | --- | --- | --- | --- | --- |
| cisterna \ vesicle | C1 | C2 | C3 | C4 | C5 |
| C1 | 0 | 1 | 0 | 0 | 0 |
| C2 | 0 | 0 | 1 | 0 | 0 |
| C3 | 0 | 0 | 0 | 1 | 0 |
| C4 | 0 | 0 | 0 | 0 | 1 |
| C5 | 0 | 0 | 0 | 0 | 0 |
| vesicular transport |  |  |  |  |  |
| cisterna \ vesicle | C1 | C2 | C3 | C4 | C5 |
| C1 | 0, 0.5 or 1 | BW 1 FW 0 | 0 | 0 | 0 |
| C2 | BW 0 FW 1 | 0, 0.5 or 1 | BW 1 FW 0 | 0 | 0 |
| C3 | 0 | BW 0 FW 1 | 0, 0.5 or 1 | BW 1 FW 0 | 0 |
| C4 | 0 | 0 | BW 0 FW 1 | 0, 0.5 or 1 | BW 1 FW 0 |
| C5 | 0 | 0 | 0 | BW 0 FW 1 | 0, 0.5 or 1 |
| distillation transport |  |  |  |  |  |
| cisterna \ vesicle* | C1 | C2 | C3 | C4 | C5 |
| C1 | 1 | 0.1 | 0 | 0 | 0 |
| C2 | 0.1 | 1 | 0.1 | 0 | 0 |
| C3 | 0 | 0.1 | 1 | 0.1 | 0 |
| C4 | 0 | 0 | 0.1 | 1 | 0.1 |
| C5 | 0 | 0 | 0 | 0.1 | 1 |

\*two cisternae carrying the same domain could also fuse

As parameters of the Golgi stability, the relative cisterna area for each Golgi domain was plotted throughout the simulation (Fig. 1C). An even distribution of areas would indicate a well-balanced Golgi. This distribution would have a maximal Shannon entropy (each domain occupying 20% of the total cisternae renders an inter-cisternae entropy = 1.6). Also, in ideal conditions, individual organelles would carry a single Golgi domain; this distribution would render a minimal internal entropy (for example, if 100% of the area of a cisterna is a C1 domain, its intra-cisterna entropy is 0). These parameters are plotted throughout the simulation (see Methods for more details). The progression of this model showed an unstable Golgi, with not all cisternae present at all times (not shown). To stabilize the Golgi, a simple solution was to duplicate the C1 initial cisterna. With this initial setting, the Golgi structures were stable and the initial fluctuations were attenuated converging to a periodicity dictated by maturation with high inter- and low intra-cisterna entropy.

The stability of this Golgi model depends on the balance between the maturation and the rate of vesicle budding. At a high fission rate, the Golgi became vesiculated and disorganized. Too low fission probability made maturation the prevailing process and Golgi resident proteins were lost (in Fig. 1C, fission probabilities of 0.002, 0.01 and 0.1 are shown).

To assess the transport capability of this Golgi, two different cargoes were included in the C1 cistern: a large soluble cargo and a small soluble cargo. The large cargo could not enter into vesicles and it was retained in the cisternae. The small cargo instead distributed during fission according to the volume of the two structures formed. To mimic a Golgi resident enzyme, a membrane-bound cargo was also included. This cargo could not enter into vesicles except in the C3 cisternae where it was recruit into the C3-formed vesicles (this would mimic the retrograde transport of a medial Golgi resident enzyme).

To measure transport, the simulation calculated the amount of each cargo associated with the different Golgi domains (C1 to C5). As expected, the large cargo was transported at the rate of the maturation of the cisternae (Fig. 1D, top middle panel). In contrast, the small soluble cargo, which could diffuse into all vesicles, was delayed in the Golgi and exited with exponential decay kinetics. This was more evident when the vesicle budding rate was high (Fig. 1D, top right panel). The Golgi-resident enzyme was efficiently retrieved from the C3 cisterna by vesicles and remained cycling between C2-C3 cisternae for extended periods of time (Fig. 1D, bottom left panel). Notice, however, that at low fission probability (p = 0.002), the Golgi resident enzyme could not be retrieved and was lost by maturation (Fig. 1D, bottom middle panel). Also, when the amount loaded in the Golgi was large, the vesicle capacity for backward transport was saturated and the enzyme was transported out of the Golgi (Fig. 1D, bottom right panel).

To assess whether the transport depends on the initial conditions, a second wave of transport was set at tick 30.000 (the newly formed C1 cisterna at this tick was loaded with cargoes). Notice that transport was very robust and occurred efficiently even under conditions where the Golgi was not stabilized (there were no major differences between the transport after the first and the second uptake).

### Vesicular Transport Model

Vesicular Transport states that cargoes coming from the ERGIC are transported by vesicles that are formed in the different Golgi cisternae and fuse with the following one in the cis-to-trans direction. Conversely, another set of vesicles move in the trans-to-cis direction carrying backward cargoes. This simple description is captured in the cartoon shown in Fig. 2A, which is a modification of the one shown in (Alberts *et al*., 2015). According to this representation of the Golgi transport, the process can be described by the following rules:

**Figure 2.**
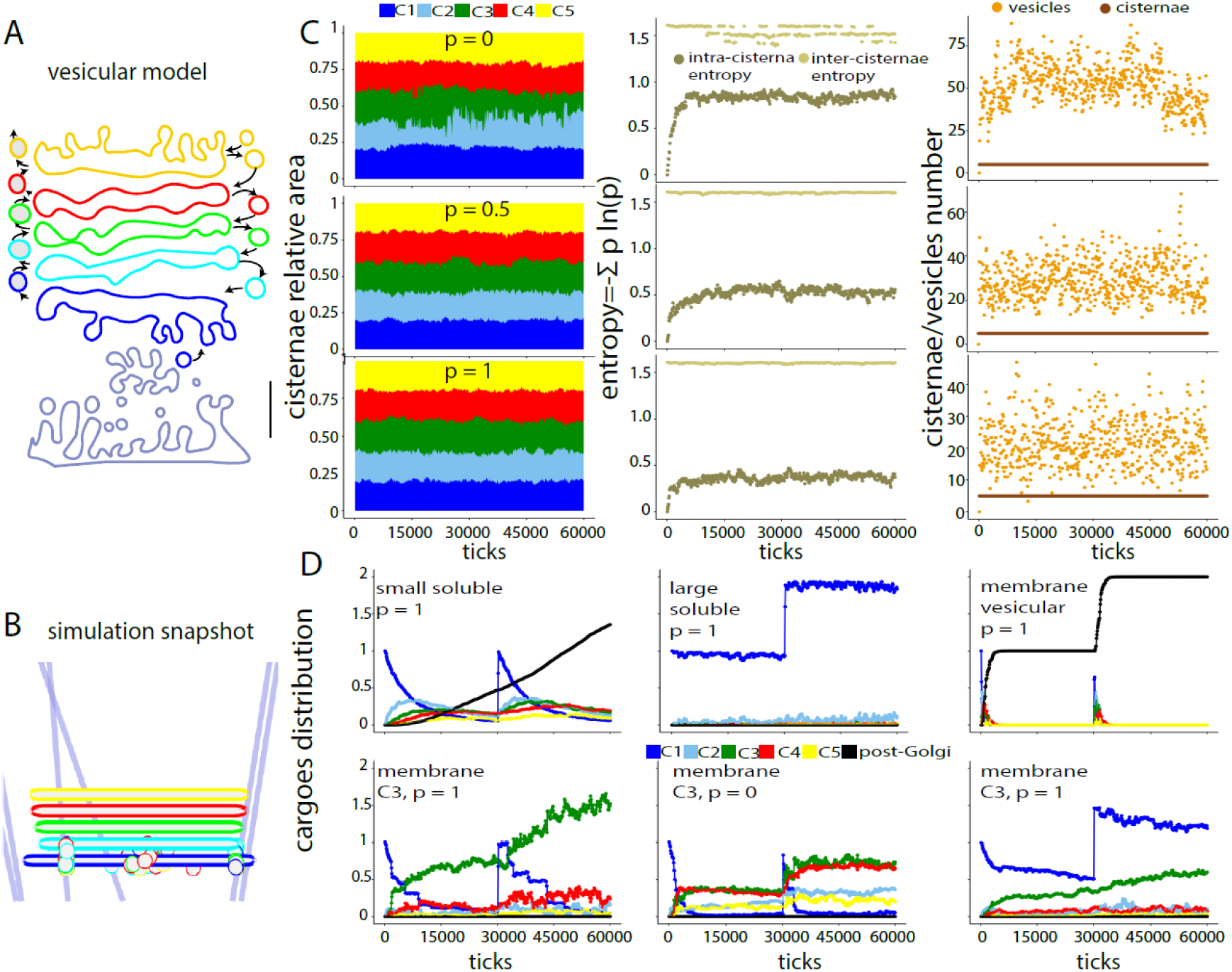
Vesicular Model. **A**. Cartoon representing the vesicular transport model. Notice that two types of vesicles can form from the cisternae. One type carries cargoes and fuses only with the following cisterna in the cis-trans direction. The other type fuses with the previous cisterna. **B**. Snapshot of the simulation built from the cartoon in A. **C**. Golgi stability when the homotypic vesicle-cisterna fusion probability was increased from 0 to 0.5, or 1. The parameters along the simulation shown are: the proportional cisterna area for each Golgi domain (left panels), the inter-cisternae and intra-cisterna entropies (central panels), and the number of vesicles and cisternae in the simulation (right panels). **D**. Transport of several cargoes under different homotypic fusion probabilities (p). The cargoes (0.004 mM) were loaded at tick 1 and 30.000. The association of a small and a large (that cannot fit into vesicles) soluble cargoes with the different Golgi domains or a post-Golgi compartment was followed throughout the simulation. Notice that the small soluble cargo was transported efficiently (top left panel), but a large cargo was retained in the C1 cisterna (top middle panel). A membrane cargo that was recruited into vesicles was rapidly transported (top right panel). A Golgi resident membrane cargo with affinity for C3 was transported to the C3 cisterna (bottom left panel). However, transport was less efficient when homotypic fusion was prevented (bottom middle panel) or when the amount of the cargo was increased a hundred times (0.4 mM, bottom right panel). In D, the results are normalized considering the maximal amount of cargo present in the simulation.

- All cisternae form two different types of vesicles.
- Forward (FW) vesicles bud from one cisterna and can only fuse with the following cisterna in the cis-to-trans direction.
- Backward (BW) vesicles bud from one cisterna and can only fuse with the following cisterna in the trans-to-cis direction.
- Forward vesicles carry cargoes that move forward in the secretory pathway.
- Backward vesicles are empty or carry cargoes moving backward in the secretory pathway.
- Vesicles are not allowed to fuse among them.
- Cisternae are not allowed to fuse among them.

To implement these rules in the ABM model, five cisternae carrying the C1 to C5 domains were generated. The cisternae could form 35 nm vesicles of two different kinds (FW and BW). The FW vesicles could only fuse with the following cisterna, whereas the BW vesicles could only fuse with the previous cisterna (see fusion probability Table I). BW vesicles forming from the C1 cisterna were deleted (they leave the Golgi to fuse with ER/ERGIC elements). FW vesicles forming from the C5 cisterna also were deleted since they fuse with the TGN. A snapshot of the simulation is shown in Fig. 2B.

The progression of this model showed that the Golgi rapidly lost the C1 and C5 cisternae (not shown). Clearly, the model required the incorporation of these domains coming from the TGN and ERGIC as depicted in the cartoon as vesicles moving toward the cis and trans cisternae. To implement this, a decrease of the C1 (or C5) area triggered the incorporation of the equivalent of a vesicle with a C1 (or C5) membrane domains to the C1 (or C5) cisterna. With this inward flux of membrane compensating the outward flux, the Golgi was rather stable.

However, the parameters for the Golgi stability were not as good as for the maturation model. The five cisternae were not always present and there was a mixture of domains in each cisterna (reflected in low inter-cisternae entropy and high intra-cisterna entropy, Fig. 2C). An improved Golgi structure was obtained by allowing homotypic fusion of vesicles with the corresponding cisterna. Notice the better distribution of the Golgi area among the five cisternae (high inter-cisternae entropy) and the decrease in the intra-cisterna entropy (indicating less mixing of domains in each cisterna) when the probability of homotypic fusion was increased from 0 to 0.5 or 1 (Fig. 2C).

When cargoes were included in C1, as expected, the large one (that cannot enter into vesicles) was not transported and remained in C1 (Fig. 2D top middle panel), whereas the small cargo was efficiently transported from C1 to C5 and eventually left the Golgi (Fig. 2D top left panel). The transport rate of the cargo depended on the possibility of being packed in vesicles. A membrane-associated cargo that was preferentially recruited in FW vesicles was rapidly transported (Fig. 1D, top right panel). A Golgi resident enzyme was modeled as a cargo with affinity for the C3 domain and it was efficiently retained in the C3 cisterna (Fig. 2D bottom left panel). Decreasing the homotypic fusion probability (Fig. 2D bottom middle panel) or increasing the concentration of this cargo during uptakes (Fig. 2D bottom right panel) caused a defect on the transport to the C3 cisterna.

### Iterative Fractionation (Distillation) Model

“… sorting may be accomplished in a more continuous fashion by many iterations of a sorting step. The sorting step need not be particularly efficient since, like a fractional distillation apparatus, high efficiency sorting would result from repetition of the sorting step.”(Dunn *et al*., 1989).

The idea that intracellular transport of different cargoes is an iterative sorting process was initially suggested for the endocytic route as a way to account for the efficient transport of ligands to lysosomes, and receptor sorting and recycling to the plasma membrane (Dunn *et al*., 1989). However, the mechanism is general enough to be applied to any transport. In brief, fusion among organelles carrying compatible membrane domains promotes the mixing of compartments whereas fission causes the separation of membrane domains preserving the identity of the compartments. Cargoes in the interior of these organelles have then the possibility of interacting with different membrane domains and during fission, they are sorted according to the affinity for these membrane structures. Cargos without any affinity for membrane domains, distribute homogeneously in the new organelles formed by fission. This model adapted to the Golgi structure is represented in the cartoon of Fig. 3A and can be described by the following rules:

**Figure 3.**
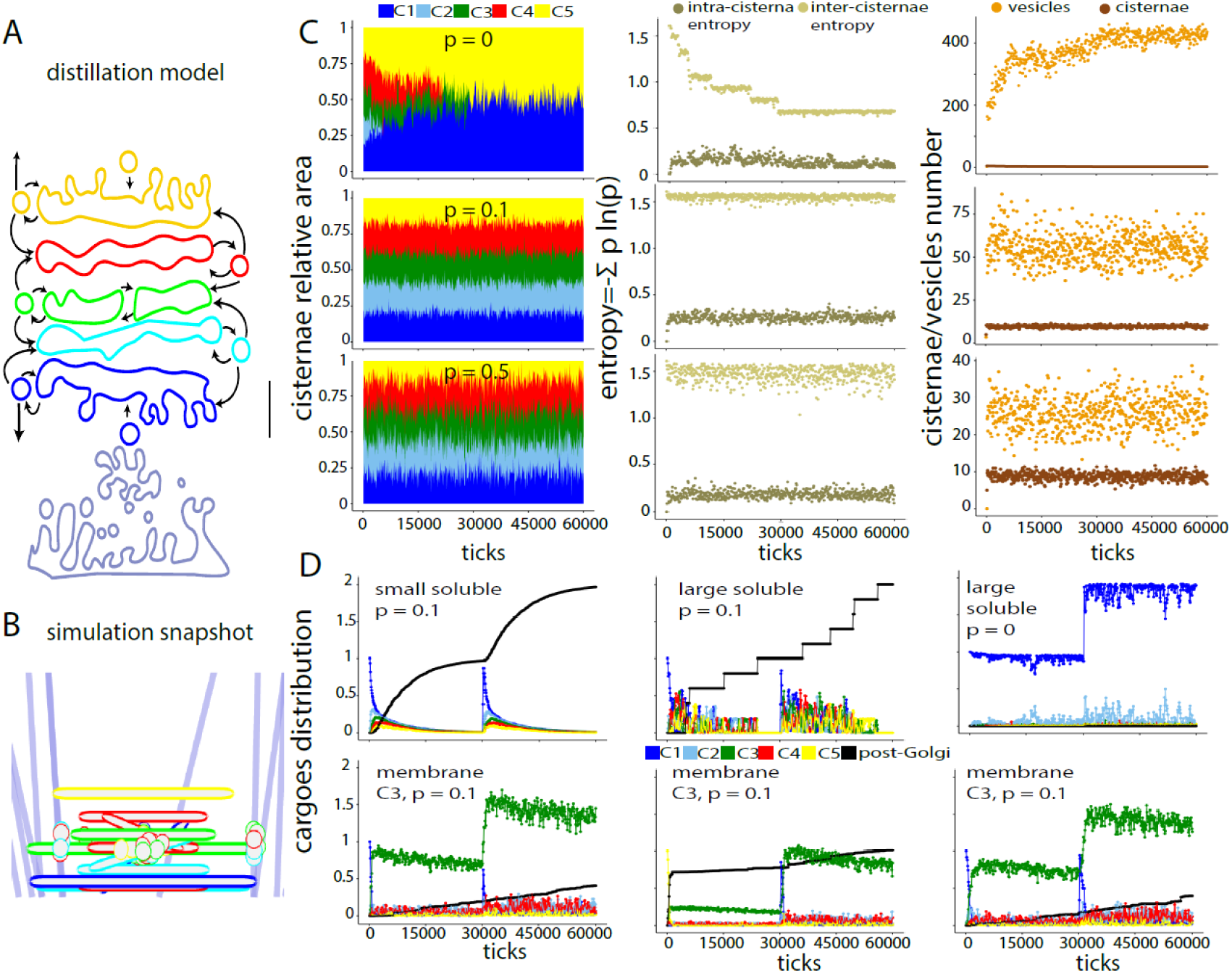
Distillation Model. **A**. Cartoon representing the distillation model. Notice the possibility of undergoing homo and heterotypic fusion. Homotypic fusion among cisternae is also allowed. **B**. Snapshot of the simulation built from the cartoon in A. **C**. Golgi stability when the probability of forming a single-domain cisterna was increased from 0 to 0.1 or 0.5. The parameters along the simulation shown are: the proportional cisterna area for each Golgi domain (left panels), the inter-cisternae and intra-cisterna entropies (central panels), and the number of vesicles and cisternae in the simulation (right panels). **D**. Transport of several cargoes under different single-domain cisterna formation probabilities (p). The cargoes (0.004 mM) were loaded at tick 1 and 30.000. The association of a small and a large (that cannot fit into vesicles) soluble cargoes with the different Golgi domains or a post-Golgi compartment was followed throughout the simulation (top panels). Notice that on the right panel, the cisterna-formation probability was 0 and the large cargo was not transported. In the bottom panels, the same association was plotted for a membrane-bound cargo with affinity for C3. Notice that, in the middle panel, the first pulse of cargo was loaded in C5 and the second in C1. A large proportion of the first pulse left the Golgi due to the membrane flux; however, the rest was correctly delivered backward to the C3 compartment. The right panel shows the transport when the cargo concentration was increased to 0.4 mM. In D, the results are normalized considering the maximal amount of cargo present in the simulation.

- All cisternae form a single type of vesicle carrying forward and backward cargoes.
- Vesicle budding from a cisterna can fuse with the same cisterna (homotypic fusion) or with the following or preceding cisternae (heterotypic fusion).
- Vesicles are not allowed to fuse among them.
- Only cisternae carrying the same membrane domain can fuse.

To implement these rules in the ABM model, the same five cisternae carrying the C1 to C5 domains were generated. The cisternae could form 35 nm radius vesicles surrounded by C1-C5 domains. The vesicles could fuse according to the fusion compatibility shown in Table I (high probability of fusing with its own cisterna, lower probability of fusing with the preceding or following cisterna and null probability of fusion with other cisternae). Vesicles forming from the C1 cisterna were deleted randomly (mimicking the fusion with ER/ERGIC elements). To prevent retrograde transport to the ERGIC, the content of the deleted C1 vesicles were transferred to the largest cisterna carrying C1. Structures carrying C5 could also be deleted mimicking transport to the TGN. The probability of being selected for deletion was inversely proportional to the area of the C5 structure (large C5 organelles have less probability of being deleted). As for the vesicular transport model, new C1 and C5 domains were incorporated into the systems triggered by a decrease of the C1 or C5 areas. Membrane domains, volume, and area of the cisternae and vesicles were preserved during the fusion and fission steps. A snapshot of the simulation is shown in Fig. 3B.

The progression of this model showed that the Golgi was unstable with a poor separation among cisternae (Fig. 3C, top panels). The high flexibility for fusion between vesicles of different origin with a cisterna promotes the mixing of Golgi domains. In the iterative fractionation model, fission is important to maintain the separation among membrane domains. This was evident in the endocytic pathway simulations (Mayorga and Campoy, 2010); compartments maintained their identity by forming not only vesicles but also large tubules. Similarly, the Golgi recovered its structure when the budding of a membrane domain was not restricted to form a 35 nm vesicle and was extended to larger cisterna-like structures. Fig. 3C, middle panel shows the Golgi stability when the probability of budding structures larger than a vesicle was increased from 0 to 0.1. Notice that better parameters were obtained with the 0.1 probability; beyond this value, the inter-cisternae entropy decreased (Fig. 3C, bottom panel).

In this model, transport was also efficient. Membrane cargoes distributed according to their affinity for the different Golgi domains. Soluble cargoes with no affinity for Golgi domains distributed according to the volume of the cisternae. In the simulation implemented here -with all cisternae having similar volume and area-the destination of this cargo depended on the membrane flux generated by the disappearance of vesicles and cisternae at the trans side of the Golgi. Notice that a large soluble cargo was not transported unless the budding of structures larger than a vesicle was allowed (Fig. 3D, top right panel). Cargoes with affinity for a specific Golgi domain could travel forward or backward to find its target. A cargo with affinity for the C3 domain loaded in C1 was rapidly transported to the C3 cisterna and remain there for extended periods of time (Fig. 3D, bottom left panel). When loaded in the trans side, part was secreted since retrieval from post-Golgi structures was not implemented in the model, but the rest was directed to the C3 cisterna (Fig. 3D bottom middle panel). The distribution of this cargo was resistant to a hundred increase of its concentration (Fig. 3D bottom right panel).

### Adding Glycosylation to the Distillation Model

As an example of this strategy, a set of cisternae containing three glycosylation enzymes (E1, E2, E3 with affinity for C1, C2, and C3 Golgi domains, respectively) were allowed to stabilize for 30.000 ticks (about 30 min, see Methods for this equivalence). Then, a membrane-bound substrate for these enzymes was loaded in a C1 cisterna at 0.01 mM concentration and the glycosylation of the substrate as it traveled through the Golgi was followed for another 30 min. A cartoon of the model is shown in Fig. 4A. The reactions implemented in COPASI are listed in Table III. Notice that the enzymes were conveniently localized in the cisternae as the cargoes were transported and glycosylated (Fig. 4B bottom panels). The kinetics of the enzymes was adjusted to prevent that a significant amount of substrate left the Golgi only partially glycosylated (most of the molecules recovered in post-Golgi structures were fully glycosylated; Fig. 4B right panel on the middle). Repast allows following the glycosylation reactions in any of the organelles. As an example, the glycosylation is shown in two vesicles and three cisternae after 200 ticks in Fig. 4C.

**Table III.**
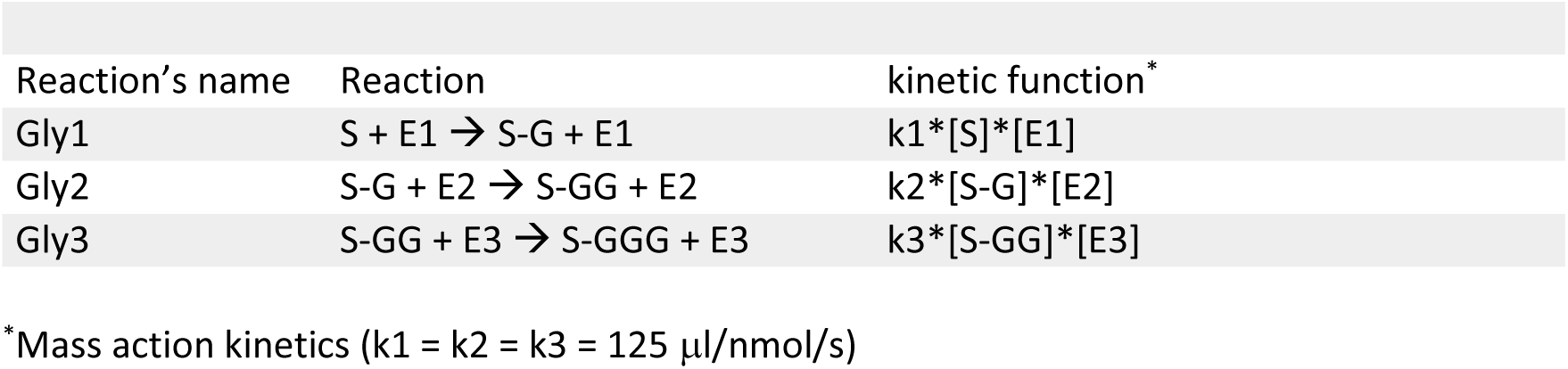
Set of irreversible reactions and kinetic functions programmed in COPASI.

**Figure 4.**
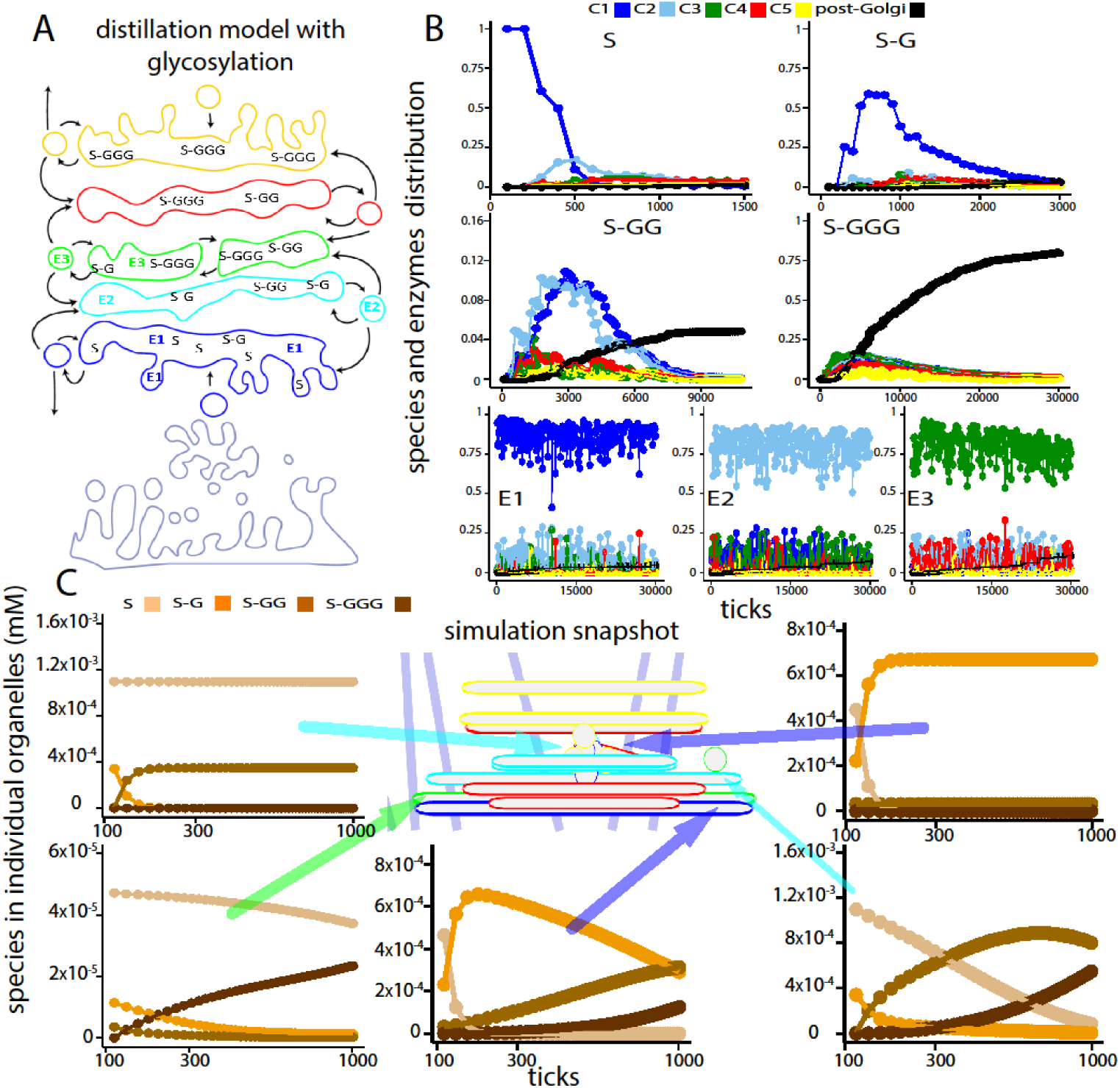
Glycosylation Simulated within the Distillation Model. **A**. Cartoon representing the distillation model where a substrate is glycosylated by three enzymes (see reactions in Table III). In the distillation model, vesicles carry both cargoes and enzymes. **B**. Transport of the unglycosylated and glycosylated species (S, S-G, S-GG, and S-GGG), and three glycosyltransferases (E1, E2, and E3, with affinity for C1, C2, and C3 domains, respectively). The enzymes (0.004 mM) were equilibrated for 30000 ticks and then S was included in a C1 cisterna at 0.01 mM concentration. The association of the species and enzymes with the different Golgi domains or a post-Golgi compartment was followed throughout the simulation. In B, the results are normalized considering the maximal amount of cargo present in the simulation. **C**. The simulation was stopped at tick 200 and the glycosylation time series calculated by COPASI for five individual organelles (two vesicles, top panels, and three cisternae, bottom panels) were plotted. The arrows point to the five organelles analyzed. The color of the arrows indicate the prevailing Golgi domain of the organelles (blue, C1; cyan, C2; green, C3)

Molecular interactions and chemical reactions can easily be implemented on top of the ABM model (Mei *et al*., 2014). Each agent can send its composition to the ODE-solving software COPASI that will calculate the molecular changes according to a series of differential equations. COPASI will return a time series with these changes that will be used to update the composition of the organelle. COPASI works with physical time and molecular units (moles or number of molecules). Repast with ticks, area units (for membrane-associated molecules), and volume units (for soluble molecules). The conversion we have applied is explained in Methods. Whenever an organelle undergoes a change due to an ABM action (e.g., fusion or fission), COPASI is called and a new time series is calculated.

## DISCUSSION

Intracellular transport is a very dynamic process involving organelles that move actively, changing shape and composition, and that undergo dramatic rearrangements of membrane and soluble factors by fusion with other organelles and budding of tubules and vesicles. It is hard to put together all these events to formulate a hypothesis about the underlying logic of transport of lipids, proteins, and carbohydrates. However, for years it has been evident from a large set of experimental approaches that membrane and soluble factors of different nature find their ways into the labyrinth of intracellular compartments in a robust and predictable way. Moreover, hundreds of factors required for the process have been identified and their function carefully characterized; many of them have been related to human diseases (De Matteis and Luini, 2011).

Despite of all these data, the mechanisms are still not well understood. In books, reviews and publications, compartments are depicted as static structures and transport is represented by arrows, frequently missing the dynamic changes observed in real-time movies.

In this report, we want to stress the necessity of more realistic models that capture the essence of intracellular trafficking and also to provide a modeling strategy that is flexible enough to translate a schematic drawing into a functional simulation that is able to generate quantitative predictions.

As an example of the flexibility of this modeling strategy, we have simulated the maturation and vesicular hypotheses for Golgi transport in their more classical and simplistic versions (Alberts *et al*., 2015). We also modeled the iterative fractionation or distillation transport mechanism that we have previously used for the endocytic route, adapted to the characteristics of the Golgi apparatus (Mayorga and Campoy, 2010).

It is important to stress that the different models for Golgi transport share several common features. The three requires fusion of membrane-bound structures and fission of budding vesicles/cisternae. These two processes are key for intracellular trafficking. Another common feature is membrane flux. The Golgi apparatus is not a closed system and requires the influx and efflux of membrane-bound structures. This is especially evident for the maturation model, but it is also present in the vesicular transport and distillation models.

Maturation, at present the most widely accepted model, needs to postulate additional mechanisms to fit the experimental data. For example, the fact that albumin (and other small soluble proteins) travel through the Golgi faster than procollagen (a large cargo that cannot enter into vesicles) does not fit with the model (Fig. 1D, top panels). To account for these observations, dynamic connections among cisternae are postulated (Beznoussenko *et al*., 2014). These connections would permit the fast diffusion of small cargoes. On the other hand, vesicular transport also requires additional transport mechanisms. By itself, it cannot account for the efficient transport of large cargoes that cannot fit into vesicles (Fig. 2D top middle panel). So, the compartment carrying these cargoes are postulated to change by sporadic heterotypic fusion with adjacent stacks that would allow the transport of large cargoes without leaving the cisternae (Lavieu *et al*., 2014). The iterative fractionation model has several attractive features. It applies to the endocytic and secretory pathways and it can be interpreted as a version of the maturation and vesicular transport models. In yeast, the switch of a cargo-containing Golgi compartment by the acquisition of a trans marker as the cis marker was leaving has been well documented (Kurokawa *et al*., 2019). In the report by Nakano’s group, both markers were membrane proteins, so the switch could not be done by exchanging factors with the cytosol and required the incoming and outgoing of membrane-bound organelles. In the distillation model, vesicles are not only carriers for cargo molecules; they are also vehicles for the transfer of membrane domains, and hence they can actively participate in the switch of cisterna identity. Notice that as postulated by the maturation model, they carry Golgi resident molecules (Fig. 4A). The distillation model has also some common features with vesicular transport assuming a single set of vesicles that transport not only cargoes but also Golgi domains. Forward or backward transport is dictated by the affinity of cargoes for different Golgi domains. Soluble and membrane-associated molecules with no specific affinity, would be transported by the flux of material that is added at the cis side of the Golgi and is withdrawn at the trans side.

In this report, we show that the basic rules that support a hypothetical transport mechanism can be extracted from a schematic drawing, and that these rules are sufficient to build an ABM simulation rendering quantitative predictions. It is important to stress that each rule should have an underlying molecular mechanism that we have not explored. For example, the hypothesis that vesicles budding from a cisterna fuse only with the preceding (or the following) cisterna would require the identification of factors involved in specific recognition and fusion and the maintenance of these factors in the correct localization to be incorporated into vesicles for the next round of fusion. In this sense, the distillation model has a simple explanation for Golgi homeostasis. According to this model, the Golgi domains are stable structures budding vesicles that fuse predominantly in a homotypic way; heterotypic fusion is allowed only with neighbor cisternae. Patches of membrane domains which do not correspond to a specific Golgi compartment will be incorporated into budding vesicles that will preferentially fuse homotypically, restoring the factors to their original compartment.

ABM implemented in Repast has the advantage of being compatible with COPASI, a very well-established software to handle ODE (Hoops *et al*., 2006). Therefore, in the skeleton of dynamic organelles, complex networks of molecular interactions and chemical reactions can be included. This makes the modeling suitable for many processes that heavily depends on intracellular trafficking, such as receptor signaling, antigen processing, and cellular infections. As a very naïve example, the glycosylation of a hypothetical factor by three different enzymes located to different cisternae of the Golgi was implemented in COPASI. In this simulation, glycosylation occurred in dynamic structures that continuously change composition as the glycosylated factor and the enzymes are transported through the Golgi.

It is clear that our approach is not the first mathematical model implemented to represent the Golgi structure and the transport of cargoes (Vagne and Sens, 2018a;Vagne and Sens, 2018b;Dmitrieff *et al*., 2013;Binder *et al*., 2009;Ispolatov and Musch, 2013;Mukherji and O’Shea, 2014;Sachdeva *et al*., 2016;Gong *et al*., 2010). Several research groups have published different models addressing organelle self-organization and protein and lipid sorting in the Golgi (for a review, see (Sens and Rao, 2013). Fusion, fission, and maturation are at the core of most of these models. They are based on physical principles with different degree of mathematical complexity and most can be analytically solved. Our method is in these respects modest. Its advantages are its simplicity and flexibility that would be crucial for building more complex pathways incorporating organelles of different nature. It is also easier to connect with cell biologists’ hypotheses. The schematic drawings of compartments connected with arrows can be conveniently represented in ABM and the molecular interactions in ODE, providing a multiscale support to the simulations. This modeling strategy can be used to address issues directly linked to the mechanism of transport (e.g., Rab dynamics) or as a way to incorporate the complexity of transport to other cellular processes that occur in dynamic organelles (e.g., antigen presentation or cell infection).

All considering, the goal of this report was to show that dynamic models can be built extracting the rules implicitly present in Golgi transport cartoons. At present, rules need to be programmed in Java; we have not generated a complete set of rules to choose from. We offer to help in the building of any model that interested groups may require. A long term goal would be to make accessible more user-friendly tools to recreate a complete set of rules and to extend the model to embrace the endocytic and secretory pathways in a single simulation.

## Supporting information

Movie-Distillation

Movie-Maturation

Movie-Vesicular

## METHODS

### Agent-Based Model (Repast)

The freely available modeling platform Repast (North *et al*., 2013) was used to model agents and actions in an Eclipse environment (https://repast.github.io/). The code can be accessed from the Git repository https://github.com/ihem-institute/immunity

### Ordinary Differential Equations (COPASI)

ODEs were programmed in COPASI (Hoops *et al*., 2006)(http://COPASI.org/). All COPASI files are included in the Git repository. COPASI and Repast interaction is achieved as described before (Mei *et al*., 2014). Basically, Repast sends initial concentrations present in each organelle to COPASI that generates a time series. A matrix with time series for each metabolite is sent back to Repast.

### World

The space represented is a projection in 2D of a cytosol square of 4.5 × 4.5 μm. The upper border corresponds to the plasma membrane and the lower border to the nucleus. The right and left borders form a continuous. Hence, the world shape corresponds to the surface of a cylinder.

### Time

The tick duration was calibrated with the fastest process in the model (movement of organelles on microtubules: 1 μm/sec). In the simulation, an agent requires 75 ticks to travel 4.5 μm when associated with a microtubule; hence, one tick corresponds to about 0.06 sec. The time for all other actions was adjusted changing the probability of being performed at each tick. Actions occurred every 0.06 seconds with the probabilities shown in Table I.

### Cisternae and vesicles

Each Golgi structure (cisterna or vesicle) has area and volume. The area is occupied by one or more of five Golgi domains (C1 to C5). The structures also carry membrane and soluble cargoes. In Repast, soluble cargoes were expressed as a fraction of the organelle volume, and membrane-associated cargo as a fraction of the organelle area. We assumed that these fractions roughly correspond to concentrations in mM units. According to this assumption, about 20 molecules of a soluble cargo at 1 mM concentration will be present in a 20 nm radius vesicle. This value fits well with the reported range of membrane proteins in an average synaptic vesicle of 21 nm radius (2-70 molecules, (Takamori *et al*., 2006)). The cargoes were loaded at 0.004 or 0.4 area or volume ration (corresponding to 0.004 or 0.4 mM, respectively). The transport capacity of an organelle of soluble or membrane cargoes was limited to 1 mM. No cargo was allowed to exceed this concentration making transport a saturable process.

The shape and size of cisternae correspond to 20 nm high cylinders with the area and volume of the cisterna. The cylinders were represented as round-corner rectangles in the 2D representation of the world. Vesicles are 35 nm radius spheres and are shown as circles. Cisternae and vesicles can perform the following actions:

### Move

When near microtubules (light blue straight lines in the model), vesicles and small cisternae move to the minus end of the filament (toward the nucleus). Otherwise, vesicles move randomly. Large cisternae (>250 nm radius) move randomly parallel to the nucleus in a restricted area (centered on the World and up to 360 nm from the nucleus).

### Fusion

Vesicles and cisternae sensed all other structures at a distance less than its size (the radius of a sphere with the organelle’s volume). If nearby structures carry a compatible membrane domain, they fuse. Compatibility was calculated as previously described (Mayorga and Campoy, 2010). The probability of fusion of single domain structures is specified in Table II for the different models. For structures carrying more than one domain, the probability was adjusted according to the proportional area occupied by each membrane domain. After fusion, a single organelle was formed carrying the area and volume and all the membrane and soluble components of the original structures.

### Fission

Cisternae have enough membrane to bud vesicles or another cistern. Fission always generates vesicles/cisternae carrying a single membrane domain. The domain that was incorporated in the new organelle was selected at random. The probability of budding vesicles/cisternae was proportional to the area of the cisterna. The probability was set to p = (organelle area – area of a 250 nm radius cisterna)/ (area of a 500 nm radius cisterna– area of a 250 nm radius cisterna). Soluble content distributed proportionally to the volume of the formed structures. Membrane cargo distributed proportionally to the area of the two new organelles except when they have affinity for specific Golgi domains. In this case, they were directed to the new structure if they have more affinity for the Golgi domain forming the vesicle/cisterna than for the Golgi domains remaining in the cistern. Large cargoes could not enter into vesicles and remain always in the cisternae. Soluble and membrane-bound cargoes occupied volume and area of the structures; hence, the budding structures carried, at most, the cargoes corresponding to the vesicle/cisterna volume or area. The area, volume, membrane, and soluble contents were preserved during fusion and fission events. Golgi domains were also maintained, except during the “maturation” (see specifications for this action).

### Outflux

C1 or C5 vesicles and cisternae had the possibility of leaving the system and were deleted. For maturation transport, the C5 cisterna left the Golgi every 3000 ticks; for vesicular transport, only C5 vesicles carrying cargo were transported out of the Golgi; for distillation, C5 structures were selected at random to leave the system. Larger structures had a lower probability of being selected. The probability was set to p =1 - (structure area – area of a 35 nm radius vesicles)/ 0.8E6 nm. Where 0.8E6 nm is twice the area of a 250 nm radius cistern. The cargoes present in the C5 structures that left the Golgi along the simulation were considered transported to a post-Golgi compartment. Vesicles budding from the C1 cisterna could also leave the system. However, the cargoes in the C1 structures leaving the Golgi were re-located in a C1 cisterna (to prevent retrograde transport in order to measure only forward transport).

### Influx

New Golgi structures were allowed to form to compensate for the domains that left the system. For maturation transport, a C1 cisterna was introduced every 3000 ticks; for vesicular and distillation transport, C1 and C5 vesicles were randomly added to the system.

### Pulse of cargoes

Cargoes were loaded in the initial C1 cistern. For distillation transport, one cargo was loaded in the C5 cisterna to show backward transport. For a second pulse, the cargoes were loaded in the newly formed C1 vesicles/cisternae at ticks 30.000.

### Microtubules

Straight lines were drawn in the model representing microtubules. In the present model, these structures can only change position with a 0.0001 probability.

### Cargo glycosylation

As an example of Repast-COPASI combination, the glycosylation of a factor by three different Golgi-resident enzymes was modeled in the distillation transport mechanism. Three enzymes (E1, E2, and E3) with affinity for different Golgi domains (C1, C2, and C3, respectively) were loaded in the model at 0.004 mM concentration. After 30.000 ticks, they arrived at a quasi-stable distribution. Then, the factor was loaded in a C1 cisterna at 0.01 mM concentration and the changes in the glycosylated species and their distribution throughout the Golgi structures were recorded for additional 30.000 ticks. In the simulation, each structure sent to COPASI its enzyme and substrate content in mM units, and received a time series with the evolution of the species along time. The series was re-calculated every time the composition of the organelle was changed by a transport event (e.g., fusion and fission). The reactions are shown in Table III.

### Model initialization

The parameters and initial organelle characteristics were loaded from a csv (comma-separated values) file generated from a spreadsheet. The COPASI file was included in the Eclipse environment to be called from Repast when needed.

Besides the graphical visualization, the model generates several output tables with data about the simulation.

### Membrane and soluble cargo distribution

The simulation calculated the amount of each soluble and membrane cargoes associated with the different Golgi domains. For example, to estimate the association of a soluble cargo with the C5 domains, the amount of cargo present in each endosome was multiplied by its relative content of C5 on the organelle (cargo content * C5 area/total area) and added to a total. As a rule, the simulations were run several times and the values plotted in the figures are the average of 3-5 runs.

### Relative area and entropies

To calculate the relative area and inter-cisternae entropy, all the cisternae of the simulation, larger than 250 nm radius, were classified according to their prevalent Golgi domain. The area of all cisternae carrying the same Golgi domain was summed and expressed as a proportion of the total cisternae area. The Shanon’s entropy for this distribution was calculated as –∑ p * ln (p), where p is the proportion for each Golgi domain. To calculate the intra-cisterna entropy, the same calculation was done for the proportions of Golgi domain areas in every single organelle in the simulation. The global intra-cisterna entropy was calculated as the area-weighted average of the organelles’ values.

## MOVIES

Representative movies of the maturation, vesicular transport, and distillation models are included as supplemental material. The color indicates the more abundant Golgi domain in each structure. Color code is the same than in the Figures. The letter S represents a single soluble small cargo. The letter M represents a large membrane-associated cargo.

## ACKNOWLEDGMENTS

This draft has not been send for publication. Any comment, suggestion or criticism that may help to improve the final manuscript will be greatly appreciated

